# Crystal structure of HERV-K envelope glycoprotein surface subunit

**DOI:** 10.1101/2025.10.24.684402

**Authors:** Nikos Nikolopoulos, Yorgo Modis

## Abstract

The most recently acquired and transcriptionally active family of human endogenous retroviruses (HERVs) is HERV-K. Of the approximately 100 copies of HERV-K in our genome, many retain the potential to proliferate by retrotransposition, express viral proteins, and form functional virus particles. Aberrant expression of the HERV-K envelope glycoprotein (Env) has been associated with cancer and neurodegeneration. Autoantibodies against HERV-K Env have been found in patients with various autoimmune diseases. Here, we report the crystal structures of the Env surface subunit (SU) from HERV-K HML-2, determined at 2.25 Å resolution. The overall fold is somewhat similar to Syncycin-2 SU and distantly related to HIV-1 gp120. The structure contains five disulfide bonds and four N-linked glycans, including one glycan that appears important for structural stability. Two extended loops form a surface for potential interactions with cell-surface receptors or other cellular factors. The structure also contains three detergent molecules, similar in structure to cholesterol, bound to hydrophobic surface patches. This crystal structure provides a platform for future studies to map autoantigenic epitopes, identify small-molecules that interfere with HERV-K activity, and extend our mechanistic understanding of retroviruses.

**IMPORTANCE:** 15% of the human genome consists of endogenous retroviruses and other virus-derived elements inherited from ancestral viral infections. Many endogenous retroviruses from the HERV-K family retain the ability to proliferate across the genome and produce virus-like particles. Aberrant expression of the HERV-K envelope glycoprotein is associated with cancer, neurodegeneration, and autoimmune disease. Here, we report the crystal structure of the HERV-K envelope glycoprotein surface subunit. The structure provides a view in atomic-level detail of the molecular components in HERV-K most likely to trigger autoimmune responses. The structure also identifies binding sites for drug-like molecules, paving the way for future studies to develop new therapeutics.

## INTRODUCTION

Endogenous retroviruses (ERVs) and other endogenous viral elements, relics of ancient germline infections, constitute approximately 15% of the human genome (1, 2). Most human ERV (HERV) loci have been rendered nonfunctional by mutations, deletions, and recombination events but HERV-K, belonging to a betaretrovirus-like group within the *Retroviridae* family and representing the most recent wave of endogenization, stands out as remarkably intact (3, 4). Many HERV-K, particularly, the human MMTV-like family 2 (HML-2) proviruses maintain full-length open reading frames (ORFs) for *gag, pol* and *env* genes (5). Notably, reconstructed consensus HERV-K genomes can produce infectious virions *in vitro* (6, 7).

The HERV-K envelope glycoprotein (Env) has been implicated in both health and disease contexts. During early embryonic development, HERV-K Env is expressed on the surface of pluripotent stem cells regulating their function via its association with CD98 and impacting neuronal differentiation (8). In adult somatic cells, HERV-K is generally repressed (9). Reactivation of HERV-K and expression of HERV-K Env is associated with a range of cancers, neurodegeneration, autoimmune diseases (10-15). Expression of HERV-K Env has been reported in melanoma cells (10), breast cancer (11), and ovarian cancer (12). Autoantibodies against HERV-K Env have been reported in patients with systemic lupus erythematosus (SLE) (14), rheumatoid arthritis (RA) (15) and amyotrophic lateral sclerosis (14).

As a functional class I fusion protein, HERV-K Env follows a canonical structure organization and is synthesized as a precursor which is cleaved into surface (SU) and transmembrane (TM) subunits (16). SU engages host factors for cell entry, heparan sulfate glycosaminoglycans serve as primary attachment factors while the CD98 heavy chain, a component of amino acid transporter LAT1, acts as a receptor (17, 18). Here we report the crystal structure of the receptor-binding unit (SU) of a human endogenous retrovirus at 2.25 Å resolution using a consensus sequence derived from ten HERV-K (HML-2) proviruses (7). Our crystal structure complements two cryo-EM reconstructions of the trimeric Env SU-TM (19, 20), published while this manuscript was in preparation, to provide a high-resolution view of the SU subunit including glycosylation and disulfide bonds. In addition, we identify detergent molecules bound within distinct SU pockets, which shed light on potential mechanisms of receptor engagement at the cell surface, SU-TM interactions and provide starting points for structure-guided therapeutic development. These insights establish an atomic framework for dissecting HERV-K Env function and exploring strategies to counteract HERV-K Env associated disease activity.

## RESULTS

### Crystal structure of HERV-K Env SU

Env SU (residues 111–430) of a consensus-sequence HERV-K construct, HERV-K_CON_ (7), was expressed in HEK293S GnTI^−^ cells to produce a uniform glycosylation pattern to facilitate glycoprotein crystallization.

HEK293S GnTI^−^ lack N-acetylglucosaminyltransferase I activity and accordingly secrete proteins with Man5GlcNAc2 (mannopentaose-di-(N-acetyl-D-glucosamine)) as the predominant N-linked glycosylation. The long N-terminal native signal peptide of HERV-K Env (residues 1–110) was replaced by the signal peptide-pro-peptide domain from human Trypsin-1 to enhance protein secretion (21). Substitution of cysteine at position 141 with alanine (C141A) abolished non-specific protein cross-linking and aggregation through aberrant disulfide bond formation. The abovementioned protein engineering procedures produced a high yield of soluble HERV-K Env SU for subsequent purification, crystallization, and crystallographic structure determination.

The crystal structure of HERV-K_CON_ Env SU was determined at 2.25 Å resolution by molecular replacement using an atomic search model generated with AlphaFold 3 (22). See **Table 1** for crystallographic data collection, refinement, and validation statistics. The fold is composed primarily of β-sheets. An extended β-sheet network at the core of the protein provides structural stability and organizes the fold (**Fig. 1A**). The SU subunit can be subdivided into an inner and an outer domain, adopting terminology from the structure of the gp120 envelope glycoprotein from HIV-1 (23, 24). The inner domain (residues 111-171 and 379-430) forms a membrane-proximal structural unit expected to interface with the transmembrane (TM) subunit, analogous to the interaction between HIV-1 gp120 and gp41 (the SU and TM subunits, respectively). The HERV-K SU inner domain consists of β-strands and loops and contains both the N- and C-termini of SU. The main structural feature of the inner domain is a distorted β-barrel of six β-strands, stabilized by three disulfide bonds, linking cysteines 164 and 171, 382 and 413, and 405 and 408. Notably, residues 405-408 are part a CXXC motif, considered a hallmark of Envs from gamma-type retroviruses. The outer domain (residues 172-378) encompasses the expected receptor-binding region at the membrane-distal apex. It contains one long β-sheet of five antiparallel β-strands and one short β-sheet of three antiparallel β-strands, separated by two long loops, each stabilized by a disulfide bond (Cys227-Cys246 and Cys275-Cys282). An α-helix is also present near the interface with the inner domain.

**Table 1.** Crystallographic data collection, refinement, and validation statistics.

| **Data collection** | HERV-K Env SU |
| --- | --- |
| Space group | *P* 1 |
| Cell dimensions |  |
| *a*, *b*, *c* (Å) | 60.04 63.52 70.68 |
| $\alpha$, $\beta$, $\gamma$ (°) | 101.94, 101.83, 104.06 |
| Resolution (Å) | 59.32 – 2.254 (2.31 – 2.25) |
| Total reflections | 162,480 (7,933) |
| Unique reflections | 44,339 (3,144) |
| Multiplicity<sup>a</sup> | 3.7 (3.5) |
| Completeness | 98.2% (97.4%) |
| $\langle I \rangle / \sigma(I)$ | 11.4 (0.7) |
| Wilson *B*-factor | 47.2 |
| *R*<sub>merge</sub> | 0.114 (1.50) |
| *R*<sub>pim</sub> | 0.070 (0.931) |
| CC1/2 | 0.979 (0.317) |
| **Refinement and Validation** |  |
| Resolution (Å) | 59 – 2.25 (2.31 – 2.25) |
| No. reflections, working set | 44,339 (3,144) |
| No. reflections, test set | 2,018 (145) |
| *R*<sub>work</sub> | 0.2265 (0.3852) |
| *R*<sub>free</sub> | 0.2864 (0.4396) |
| No. non-hydrogen atoms | 5,903 |
| Protein | 5,199 |
| Glycans | 353 |
| Ligands | 126 |
| Water | 225 |
| Average *B*-factors<sup>a</sup> |  |
| Protein (Å<sup>2</sup>) | 57.53 |
| Glycans (Å<sup>2</sup>) | 91 |
| Ligand (Å<sup>2</sup>) | 87 |
| Water (Å<sup>2</sup>) | 52 |
| Root mean square deviations |  |
| Bond lengths (Å) | 0.008 |
| Bond angles (°) | 1.13 |
| Ramachandran plot |  |
| Favored (%) | 92.23 |
| Allowed (%) | 7.29 |
| Outliers (%) | 0.48 |
| Clashscore (Phenix 1.21.1) | 14.28 |
| PDB accession code | 9SYA |

Numbers in parentheses refer to highest resolution shell.
<sup>a</sup>Residual *B*-factors after TLS refinement. See PDB entry for TLS parameters.

**Fig. 1.**
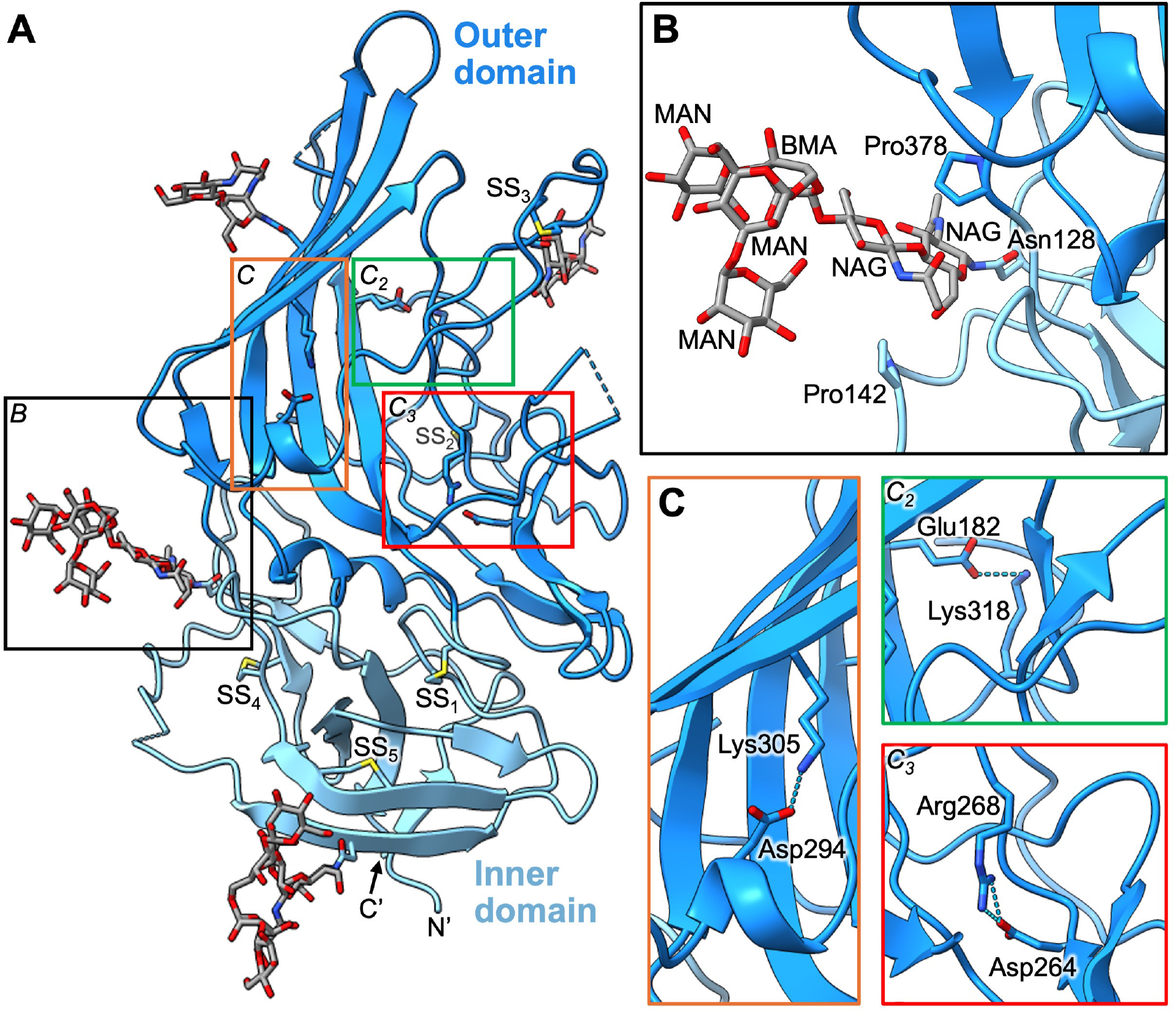
Overall fold and structural features of HERV-K Env SU. (**A**) Overall fold of (residues 111-430), consisting of inner and outer domains. Four N-linked glycans are shown in ball-and-stick representation with carbon atoms in grey. Five disulfides are shown in ball-and-stick representation with yellow sulfur atoms (SS_1-5_). The TM domain and viral membrane would be located below the N- and C-termini (labeled N’, C’). (**B**) Closeup of the hexasaccharide glycan linked to Asn128, at the interface between the inner and outer domains. NAG, N-acetyl glucosamine; MAN, α-*D*-mannose; BMA, β-*D*- mannose. See **Table 1** for crystallographic data collection, refinement, and validation statistics. (**C**) Closeups showing three salt bridges that stabilize the fold of the outer domain.

Four of the five predicted N-linked sequons (Asn128, Asn153, Asn274, and Asn355) are glycosylated in HERV-K Env SU expressed in HEK293S GnTI^−^ cells; the inner-domain sequon at Asn372 shows no evidence of glycosylation in the electron density map. At Asn128 the map contained clear density for the chitobiose core (two N-acetylglucosamines) plus four mannose moieties. Together these carbohydrates residues form a hexasaccharide that is tightly lodged in a surface cavity between the inner and outer domains, forming hydrophobic contacts with proline residues in each domain (**Fig. 1B**). The unusually well-ordered glycan at Asn128 therefore behaves more like an integral structural element (filling a void and making stabilizing contacts) than a flexible surface decoration. By contrast, the glycans at positions 274 and 355 sit on the exposed outer domain and display more limited, peripheral density consistent with solvent-accessible, mobile oligomannose adducts that are unlikely to form a continuous steric cloak. The glycan at position 153, although located in the inner domain, lacks the structure seen at position 128 but may nevertheless contribute more subtly to transient interactions with the TM subunit or modulate local folding dynamics. These observations define a spatially sparse glycan landscape on HERV-K SU that contrasts sharply with the dense glycan shield of HIV-1 gp120 (23-25) and more closely resembles the lighter glycosylation reported for Syncytin-2 SU subunit (26). The architecture implies relatively large protein surfaces will be available in HERV-K SU for receptor engagement and antibody recognition, with the glycan at Asn128 representing a localized structural exception rather than part of an immune-evasive glycan cloak.

Within the outer domain, two salt bridges play key roles in structural stabilization. The first, between Asp264 and Arg268, reinforces the short β-sheet of 3 antiparallel β-strands and its adjoining loop (**Fig. 1C**). The second, between Asp294, located within a short α-helical segment, and Lys305, stabilizes the longer five-stranded β-sheet together with the adjoining extended loop region (**Fig. 1C**). Together, these interactions delimit and stabilize the surface-exposed loop. A third salt bridge is present between Glu182 of the longer β-sheet and Lys318 of a smaller loop between the two β-sheets (**Fig. 1C**). Other notable structural features are three relatively rare cis peptide bonds involving residues Gly135-Pro136, Ser329-Pro330, and Pro385-Pro386.

### Detergent molecules bind hydrophobic surfaces at the TM interaction interface

A molecule of CHAPS (3-[(3-cholamidopropyl)dimethylammonio]-1-propanesulfonate), a cholic acid derivative included in the crystallization solution (27), was modeled within a hydrophobic pocket at the interface between the outer and inner SU domains (**Fig. 2**). The four-ring steroid nucleus of CHAPS – specifically rings C and D and one protruding methyl group on its convex face – nestles into this pocket, which is formed primarily by Phe259 (outer domain) and Pro398 (inner domain).

**Fig. 2.**
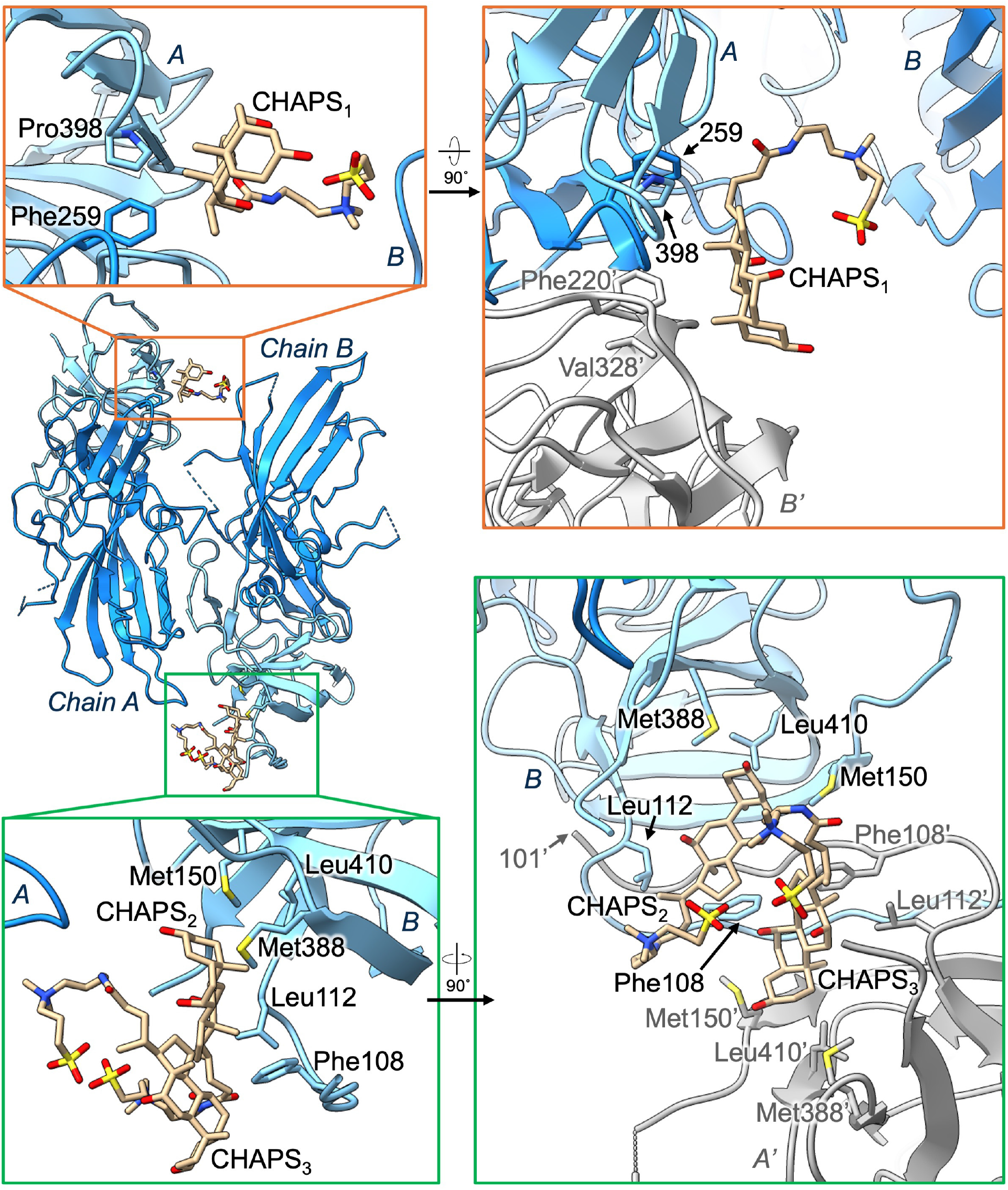
Binding mode of three sterol molecules to HERV-K SU. Three cholic acid derivative molecules were clearly identifiable in the electron density map and modeled as CHAPS (3-[(3-cholamidopropyl)dimethylammonio]-1-propanesulfonate). Cholic acid derivatives were present as additives in the crystallization condition (see **Materials and Methods**) (27). CHAPS molecules are shown in ball-and-stick representation with pale yellow carbon atoms. “*A”* and “*B”* denote two HERV-K SU chains in the crystallographic asymmetric unit. Proteins shown in grey are related to the asymmetric unit by crystallographic symmetry. Their chain and residue identifiers are show in grey and followed by a prime symbol.

Two additional CHAPS molecules were modeled on concave hydrophobic surface patches on the inner SU domain adjacent to the trypsin-derived signal-peptide tail. One of these molecules makes hydrophobic contacts with a hydrophobic surface clusters formed by Met150, Met388, Leu112, and Leu410, along with a phenylalanine from the trypsin-derived signal-peptide tail (residues 99-110; **Fig. 2**). The second CHAPS molecule is stabilized by residues from an adjacent SU molecule in the crystal lattice (crystallographic symmetry packing interactions), as well as by interactions with the neighboring CHAPS molecule. These CHAPS molecules stabilize structural features in the SU fragment that, in the intact envelope complex, are maintained via interactions with the TM subunit (19, 20).

### Comparison to other retroviral Env SU structures

A structural similarity search with DALI (28) identified Syncytin-2 SU as the protein with the most similar structure to HERV-K SU (Z-score 7.5, rmsd 5.9 Å, PDB 7OIX (26)). The next most similar structures were those of HIV-1 gp120 (Z-score 3.2, rmsd 5.4 Å, PDB 6PWU (25)) and SIV gp120 (Z-score 2.8, rmsd 3.4 Å, PDB 8DUA (29)), although the similarity scores for these were only marginally higher than for clearly unrelated proteins (Z-score 2.7 and below). Syncytin-2, which drives cell-cell fusion of trophoblasts fusion in placental development, is derived from the Env of HERV-FRD, a gammaretrovirus-like endogenous retrovirus (26, 30, 31). Most of the structural similarity between HERV-K SU, Syncytin-2 SU and HIV-1 gp120 maps to the inner domain, which has a similar distorted β-barrel as its core architectural element in all three proteins (**Fig. 3**). Upon superposition on HERV-K SU inner domain, the root mean square deviations of the inner domain Cα positions of Syncytin-2 SU and HIV-1 gp120 were 2.89 Å and 4.01 Å, respectively. Given that the inner domain is proximal to the TM membrane fusion subunit, the structural conservation of this domain suggests a common mode of intramolecular communication within the heterodimeric Env complexes.

**Fig. 3.**
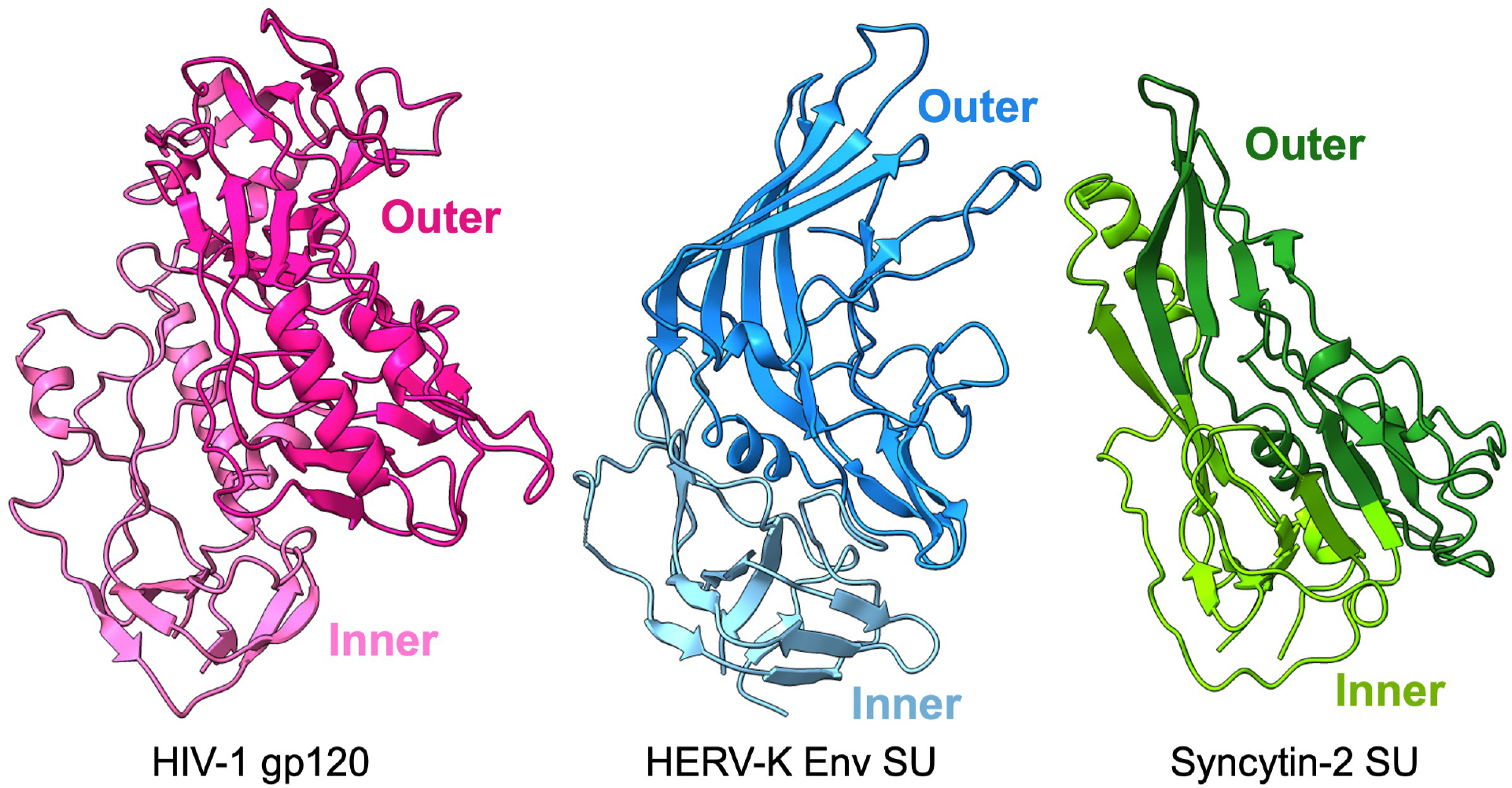
Comparison of HERV-K Env SU to Syncytin-2 SU and HIV-1 gp120. Structures of HIV-1 gp120 (PDB 6PWU; (25)), left, and Syncytin-2 SU (PDB 7OIX; (26)), right, the closest structural homologs of HERV-K SU. The relative orientations of the proteins reflect structural alignment of their inner domain using the HERV-K SU inner domain as the reference. The root mean square deviation of the inner domain Cα positions after superposition onto the HERV-K SU inner domain was 2.89 Å for Syncytin-2 SU and 4.01 Å for HIV-1 gp120.

The outer domain of HERV-K SU incorporates a mixed architecture comprising β-sheets, loop insertions, and helical elements. The main structural feature is a long β-sheet composed of five antiparallel β-strands, a topological element also found in the outer domain of Syncytin-2 SU. However, the structures of the outer domains, and the insertions within them, are otherwise highly divergent in HERV-K SU, Syncytin-2 SU and HIV-1 gp120 (**Fig. 3**).

## DISCUSSION

Here we report the crystal structure and associated atomic model of the envelope glycoprotein Surface (SU) subunit from the human endogenous betaretrovirus HERV-K (HML-2). The HERV-K SU fold retains the two-domain topology, with inner and outer domains, seen in other retroviral SU subunits including gammaretroviruses (Syncytin-2) and lentiviruses (HIV-1). This topological conservation demonstrates that a common architectural blueprint underlies Env proteins across the *Orthoretrovirinae* subfamily of *Retroviridae* (which includes the genera alpharetrovirus, betaretrovirus, gammaretrovirus, lentivirus, and others), with genus-specific refinements (glycan decorations, loop insertions, receptor specificities) overlaying the common framework. However, despite the similarity in secondary structure topology to other retroviral SUs, perhaps the most striking feature of the structure of HERV-K SU is how different its three-dimensional structure is from those of its closest structural homologs, Syncytin-2 SU and HIV-1 gp120. This indicates that HERV-K SU has diverged further from the SUs of other gamma- and gamma-like retroviruses at the structural level than might have been expected from phylogenetic analysis.

The crystal structure of HERV-K SU, determined at 2.25 Å resolution, reveals various HERV-K-specific structural elements. The fold is stabilized by five disulfide bonds and three key salt bridges. Two loop regions provide an extensive surface for potential interactions with cell-surface receptors or other cellular factors. The structure offers a detailed view of the glycan landscape. Four N-linked glycans make for a relatively sparse coverage of the molecular surface of SU, standing in contrast with the dense glycan shield of HIV-1 gp120. An unusually well-ordered, cavity-embedded branched carbohydrate chain at Asn128 of HERV-K SU appears to function as an integral structural element within the inner domain. The crystal structure also contains three amphiphilic CHAPS molecules, similar in structure to cholesterol, bound to hydrophobic surface patches of SU. These could prove to be attractive starting points for small-molecule or fragment screening campaigns to identify and compounds that bind tightly to SU and interfere with the membrane fusion activity of HERV-K Env or its ability to interact with other cellular proteins.

The HERV-K SU crystal structure reveals the conserved structural features shared with other orthoretroviral Envs while also identifying HERV-K-specific features, namely its sparse glycocalyx, structure-stabilizing glycan, hydrophobic surface patches, and extended surface-exposed loops. Autoantibodies against HERV-K Env have been reported in patients with various autoimmune diseases (14, 15). Given that SU is the receptor binding subunit in retroviruses and contains most of the known antibody binding epitopes, the epitopes of disease-causing autoantibodies against HERV-K Env are likely to map to the solvent accessible surface of SU. The structure of SU will be an invaluable tool in future studies to identify these epitopes. More broadly, the availability of a high-resolution crystal structure for HERV-K SU, along with our optimized expression construct and purification protocol, open paths toward extending our mechanistic understanding of betaretroviral envelope proteins and developing translational applications targeting HERV-K elements, the most active and recently acquired endogenous retroviruses in humans.

## MATERIALS AND METHODS

### Cloning of expression vectors

The gene encoding the SU domain of HERV-K_CON_ Env (residues 111-430) (7) was cloned into the pcDNA3.1(+) plasmid. To enhance protein secretion, an N-terminal signal peptide–pro-peptide domain derived from human Trypsin-1 was fused to the gene (21), followed by a human rhinovirus (HRV) 3C protease cleavage site. A C-terminal hexahistidine (His6) tag was added for purification purposes, preceded by a Pro-Gly sequence to facilitate efficient removal of the His6 tag by carboxypeptidase A. Additionally, the sequence was engineered to substitute cysteine at position 141 with alanine to prevent oligomerization caused by the free cysteine residue.

### Protein expression and purification

The HERV-K Env SU domain was expressed in HEK293S GnTI^−^ cells adapted for suspension culture and maintained at 37 °C in a humidified atmosphere containing 8% CO_2_ with shaking. Cells were grown in serum-free Expi293 Expression Medium (Gibco) to a final density of 3 × 10^6^ viable cells/mL with 95–99% viability before transfection. Transfection was performed using 1 mg/mL polyethylenimine (PEI) at a DNA:PEI ratio of 1:3 (w/w). Secreted protein was harvested 5 days post-transfection by collecting the supernatant after centrifugation at 300 × g for 5 min at 4 °C to pellet cells. The clarified medium was applied to Ni Sepharose Excel resin (Cytiva), washed, and eluted with 50 mM Tris pH 7.4, 300 mM NaCl containing 10 mM and 250 mM imidazole, respectively. The protein was further purified by size-exclusion chromatography using a Superdex 200 Increase 10/300 GL column (Cytiva) equilibrated in 50 mM Tris pH 7.4, 100 mM NaCl.

### Crystallographic structure determination

Crystals were grown at 18 °C by sitting-drop vapor diffusion. Purified protein was concentrated to 11.7 mg/mL, and 1 volume of seed stock was added to 5 volumes of protein solution. The final solution was mixed with an equal volume of reservoir solution optimized from the Morpheus Fusion Screen (Molecular Dimensions) (27): 32.7% PEG 8K/ethylene glycol (1:2), 0.1 M MES/imidazole pH 6.5, 90 mM Morpheus NPS, and 1.572% (w/v) Morpheus III cholic acids. Seed stock was prepared by crushing small crystals from a previous optimized crystallization experiment conducted with varying PEG 8K/ethylene glycol (1:2) concentrations (18–36%) and Morpheus III cholic acids (0.24–1.8% w/v), while maintaining constant 0.1 M MES/imidazole pH 6.5 and 90 mM Morpheus NPS. X-ray diffraction data were collected at 100 K using Pilatus3 6M and Eiger2 XE 16M detectors at Diamond Light Source beamlines I24 and I04, respectively. Data were processed with autoPROC v1.0.5 (32), and Xia2 (Dials, Aimless) (33). Molecular replacement was performed with Phaser (34) as implemented in PHENIX v1.21.1 (35) using an atomic search model of the SU domain of HERV-K_CON_ Env from AlphaFold 3 (22). An initial model was built using AutoBuild in PHENIX, manually completed with COOT v.0.9.6 (36), and iteratively refined with PHENIX. See **Table 1** for crystallographic data collection, refinement, and validation statistics.

### Structural analysis

Coordinate files were submitted to PDBsum (37) for structural feature analysis. Molecular visualization and figure generation were performed using UCSF ChimeraX (38). Structural homologs were identified using the DALI server (39), and 3D structural comparisons were carried out using the TM-align algorithm (40).

## Data availability

The atomic coordinates of the HERV-K SU glycoprotein fragment were deposited in the Protein Data Bank with accession codes 9SYA [https://doi.org/10.2210/pdb9sya/pdb].

## Acknowledgements

We thank Fabrice Gorrec (MRC-LMB Crystallization Facility) for advice on protein crystallization, and Dom Bellini at the MRC-LMB X-ray Crystallography Facility for advice in crystallographic data collection. Crystallographic data were collected on beamlines I03 and I04 at Diamond Light Source (DLS). Access to DLS was supported by the Wellcome Trust, MRC and BBSRC. This work was supported by Wellcome Trust Senior Research Fellowship 217191/Z/19/Z to Y.M. Access to Diamond Light Source was supported by the Wellcome Trust; the Medical Research Council; and UKRI (proposal number MX36838). Open access publication was funded by the University of Cambridge. To ensure open access, the University of Cambridge has applied a CC BY public copyright license to any Author Accepted Manuscript version arising.

## Notes

### Competing Interest Statement

The authors have declared no competing interest.

### Summary of Updates

The figure and table callouts in the main text have been corrected.

